# Bovine macrophages transcriptome profiling reveals divergent responses to virulent and attenuated *Babesia bovis* strains

**DOI:** 10.1101/2025.09.16.676518

**Authors:** Valenzano Magalí Nicole, Liliana Alvarez, Beatriz Valentini, Silvina Wilkowsky

**Affiliations:** Instituto de Agrobiotecnología y Biología Molecular (IABIMO), Instituto Nacional de Tecnología Agropecuaria (INTA), Consejo Nacional de Investigaciones Científicas y Tecnológicas (CONICET), de Los Reseros y Nicolás Repetto s/n, Buenos Aires, Hurlingham B1686IGC, Argentina; Laboratorio de Inmunología y Parasitología Veterinaria, EEA Rafaela, INTA, RN 34, Km 227, CC 22, 2300, Rafaela, Santa Fe, Argentina

**Keywords:** *Babesia bovis*, macrophages, RNA-seq, differentially expressed genes, virulent strain, attenuated strain

## Abstract

*Babesia bovis* is a tick-borne parasite of major economic impact in the livestock industry. Control strategies rely mainly on the use of acaricides and live attenuated vaccines. Comparative genomic analyses have shown no major differences between virulent and attenuated *B. bovis* strains, suggesting that studies on the host’s differential response may represent a key step toward clarifying the basis of disease severity and vaccine efficacy. In this study, we analyzed by RNA-seq the differential gene expression in bovine monocyte-derived macrophages after phagocytosis of erythrocytes infected with either the virulent *B. bovis* S2P strain or the attenuated R1A one. The results revealed a common transcriptional core response of several cellular processes largely centered on lymphocyte related functions, cytokine regulation, and adaptive immune signaling. In turn, the two strains elicited contrasting responses in bovine macrophages, where the virulent strain induced the enrichment of lymphocyte- and antiviral-related pathways, and the attenuated strain led to a stronger pro-inflammatory, chemotactic, and extracellular matrix remodeling signatures. Taken together, these data improved our understanding of the early transcriptional events that develop in macrophages in response to the phagocytosis of red blood cells containing contrasting *B. bovis* strains. This large dataset could be evaluated in further studies to better characterize the pathogenetic mechanisms of bovine babesiosis and to identify targets for host-directed therapeutic strategies.

## 1. Introduction

Bovine babesiosis, primarily caused by *Babesia bovis* and *B. bigemina*, is a tick-borne disease of significant veterinary and economic importance, particularly in tropical and subtropical regions where the Ixodidae tick vectors are prevalent (Jacob et al., 2020). In the mammalian host, these apicomplexan parasites invade and multiply only within bovine red blood cells. Particularly in adults, this leads to severe clinical manifestations that include high fever, hemolytic anemia, hemoglobinuria, and in many cases, cerebral babesiosis associated with sequestration of infected erythrocytes in the brain microvasculature. Of the two main species that cause bovine babesiosis, *B. bovis* infection can result in high mortality rates due to its capacity to adhere to capillary vessels and obstruct blood circulation (Bock et al., 2004; Brown et al., 2006).

The immune response to *B. bovis* involves both innate and adaptive mechanisms. The innate immune response involves the early activation of macrophages and dendritic cells, leading to the production of proinflammatory cytokines. Adaptive response is mainly mediated by opsonizing antibodies and a Th1-type cellular immune response, characterized primarily by the secretion of interferon-γ (IFN-γ) and tumour necrosis factor (TNF-α). These cytokines contribute to parasite control by enhancing splenic macrophage activation for the clearance of infected erythrocytes and by supporting the production of IgG2 (Brown et al., 2006).

Current vaccines against *B. bovis* are based on live attenuated parasites derived through serial passage in splenectomized calves or *in vitro* culture and have been used successfully in many countries (de Waal & Combrink, 2006). Vaccination with these strains induces robust and long-lasting immunity that closely mimics the protection achieved after natural infection, yet without the occurrence of severe clinical signs (Ristic, 2018). The proposed mechanism suggests that attenuation is the result of the selection of less virulent subpopulations that remain in circulation, while the more virulent parasites are sequestered in the microvasculature and thus lost during serial passages (Baravalle et al., 2012). A previous study comparing the lipid composition of virulent and attenuated *B. bovis* strains demonstrated that differences in phospholipid profiles modulate the intensity and nature of the host immune response (Gimenez et al., 2013). These findings suggest that signaling mechanisms underlying host–parasite interactions could be important determinants of disease outcome and immune activation. Therefore, understanding how the host differentially responds to virulent and attenuated strains at the molecular level is crucial to elucidating mechanisms underlying disease severity and vaccine efficacy.

Recent advances in RNA sequencing (RNA-seq) have enabled the comprehensive characterization of host transcriptional responses to apicomplexan infections (Menard et al., 2021; Terkawi et al., 2018). Moreover, transcriptomic analyses have already been successfully applied to study differential gene expression in peripheral blood mononuclear cells during *B. bigemina* infection, revealing important insights into immune activation and inflammation (Martínez-García et al., 2025). However, to date, no studies have characterized the complete bovine macrophage transcriptome in the context of the immune response induced by *B. bovis*, nor compared responses to virulent and attenuated strains.

In this work, we analyzed the host transcriptional response in bovine monocyte-derived macrophages infected with virulent and attenuated strains of *B. bovis*. By comparing the gene expression profiles induced by each strain, we aim to better understand the immune responses they trigger in the host and explore the mechanisms by which attenuated parasites elicit protective immunity without triggering severe pathology.

## 2. Materials and methods

### 2.1. B. bovis culture

The Argentinian pathogenic S2P and the attenuated R1A strains were cultured *in vitro* using a microaerophilic stationary phase system with normal bovine erythrocytes, following the method outlined by Baravalle et al., 2012. Parasites were utilized when parasitemia reached 10 %.

### 2.2. Bovine monocyte-derived macrophages (BMDM) and phagocytosis assays

BMDM obtention was performed as described in Valenzano et al., 2025. For the phagocytosis assays, 2 x 10^6^ BMDMs per well were co-cultured with red blood cells (RBCs) infected with *B. bovis* S2P and R1A strains, and with non-parasitized erythrocytes (nRBCs) as a control, during 18 h at a multiplicity of infection (MOI) of 20:1. This time point was selected to capture the early transcriptional response of macrophages following the phagocytosis of *B. bovis*-infected erythrocytes (Diotallevi et al., 2024; Ramírez et al., 2012). After infection and 3 washes with phosphate-buffered saline (PBS) buffer, BMDMs were harvested and processed. Using these experimental conditions, we found that approximately 70% of the macrophages ingested at least one RBC, confirmed by Giemsa-stained smears (Valenzano et al., 2025). All assays were performed in triplicate.

### 2.3. RNA extraction

For each replicate, cells were harvested using 1 mL of TransZol (TransGen Biotech, Beijing, China) per 6 × 10^6^ cells and 200 µL of chloroform (Sigma-Aldrich, St. Louis, MO, USA) was added. The mixture was vigorously shaken for 30 seconds and samples were centrifuged for 15 minutes. The aqueous phase was carefully transferred to a new tube, and an additional 100 µL of chloroform were added. After vigorous shaking for 30 seconds, samples were centrifuged for 10 minutes. The upper aqueous phase was recovered, and 600 µL of isopropanol (Sigma-Aldrich, St. Louis, MO, USA) and 60 µL of sodium acetate (3 M, pH 5.2; Thermo Fisher Scientific) were added. Samples were mixed by gentle inversion and incubated at −70 °C for 2 hours. RNA was pelleted by centrifugation for 20 minutes, and the supernatant was discarded. The pellet was washed with 200 µL of cold absolute ethanol (Sigma-Aldrich, St. Louis, MO, USA), followed by centrifugation for 10 minutes. After discarding the ethanol, pellets were air-dried with the tube caps open for a few minutes. Finally, RNA was resuspended in 20 µL of diethylpyrocarbonate (DEPC)-treated water (Invitrogen, Carlsbad, CA, USA). All centrifugation steps were performed at 12,000 × g for the indicated time at 4 °C. Genomic DNA present in the samples was removed with a DNase I enzyme (1 U/µg RNA; Thermo Fisher Scientific, Waltham, MA, USA) following the manufacturer’s instructions. Sample quality was assessed using the Fragment Analyzer system (Agilent Technologies, Santa Clara, CA, USA) and quantification was performed using both the NanoDrop™ spectrophotometer and the Qubit™ Fluorometer (Thermo Fisher Scientific, Waltham, MA, USA). The integrity of the RNA samples was also assessed by electrophoresis on a 1% agarose gel.

### 2.4. RNA-seq sample preparation and sequencing

Libraries were prepared using the TruSeq Stranded mRNA Library Prep Kit (Illumina, San Diego, CA, USA) starting from 2 µg of pure RNA following the manufacturer’s instructions. Sequencing was performed at the National Center for Genomics and Bioinformatics, ANLIS-Malbrán, Argentina, on a NovaSeq 6000 system (Illumina, San Diego, CA, USA) to generate paired-end 2×150 bp reads.

### 2.5. RNA-seq data analysis

The experimental design included three replicates for each condition. The quality analysis of the reads was performed using FastQC software (Andrews, 2010). The reads were pre-processed using Trimmomatic (Bolger et al., 2014) to remove the adapters. A comprehensive index was generated using the STAR aligner (Dobin et al., 2013) from the *Bos taurus* genome ARS-UCD2.0 (GCF_002263795.3), incorporating splice junction annotations with the --sjdbOverhang 100 parameter to improve alignment accuracy of eukaryotic transcripts. Sequencing reads from each sample were mapped to the reference genome using STAR (Dobin et al., 2013) with stringent parameters, allowing a maximum of 20 multiple alignments per read (--outFilterMultimapNmax 20) and were sorted by genomic coordinates (SortedByCoordinate). Transcript assembly and quantification were carried out with StringTie (Pertea et al., 2015) in reference-guided mode, applying strand-specific fragment bias correction (-rf) to generate gene-level abundance estimates. Count matrices were prepared using the prepDE.py script provided by the StringTie pipeline. Differential expression analysis was conducted using DESeq2 (Love et al., 2014). Differentially expressed genes (DEGs) were considered statistically significant if they exhibited a false discovery rate (FDR)-adjusted p-value < 0.01 with the Benjamini-Hochberg correction and an absolute log2 fold change > 1. Heatmaps and principal component analysis (PCA) were constructed in R (version 2025.05.0) using the *ggplot2* package (Wickham, 2011). Gene overlap among datasets was visualized with the BioVenn tool (Hulsen et al., 2008). Functional enrichment analyses were performed to identify overrepresented biological functions and pathways in the gene set. Gene Ontology (GO) enrichment analysis was conducted using the enrichGO function from the clusterProfiler R package to identify significantly enriched biological processes (Yu et al., 2012). Pathway enrichment analysis was carried out using enrichKEGG, from the clusterProfiler R package, to detect overrepresented metabolic and signaling pathways according to the KEGG database (Kanehisa and Subramaniam, 2002). Additionally, Reactome pathway analysis was performed using the ReactomePA package to provide complementary insights into biologically relevant pathways (Milacic et al., 2024). All analyses were conducted using default parameters, and pathways with an adjusted p-value < 0.05 were considered statistically significant. Enrichment results were visualized using ggplot2 (Wickham, 2011) generating barplots to illustrate the most significantly enriched terms and pathways.

## 3. Results

### 3.1. Sequencing

To investigate the transcriptional response of bovine macrophages upon exposure to erythrocytes infected with *Babesia bovis* strains S2P and R1A, as well as uninfected erythrocytes as a control, we generated paired-end sequencing libraries from the three experimental groups. For clarity, these conditions are hereafter referred to as S2P, R1A, and nRBC, respectively. Between 16151764 and 35361940 reads were obtained per sample (Table 1). Quality assessment indicated consistently high-quality data, with Phred scores of 35.6-35.7. The GC content ranged from 51 % to 54 % with a uniform read length of 150 bp. These results confirmed that the sequencing data were of sufficient quality for downstream analyses.

**Table 1:** Summary of read metrics and quality control results.

| Condition | Sample reference | Read count | Phred quality score | GC content (%) |
| --- | --- | --- | --- | --- |
| S2P | S2P_1 | 18958150 | 35.7 | 53 |
|  | S2P_2 | 18934108 | 35.7 | 54 |
|  | S2P_3 | 22672488 | 35.7 | 53 |
| R1A | R1A_1 | 23782786 | 35.7 | 51 |
|  | R1A_2 | 23199864 | 35.7 | 51 |
|  | R1A_3 | 16151764 | 35.7 | 51 |
| nRBC | nRBCs_1 | 35361940 | 35.7 | 51 |
|  | nRBCs_2 | 21964290 | 35.7 | 51 |
|  | nRBCs_3 | 23777810 | 35.7 | 52 |

We performed a PCA on the three replicates per condition to assess sample clustering and variability (Supplementary Data 1). One of the control replicates (nRBCs_3) showed a clear deviation from the other two and was therefore excluded. Subsequent analyses were conducted using the remaining two replicates for this condition.

### 3.2. Differential expression analyses

Across the three experimental conditions, 4679 protein-coding genes were aligned against the *Bos taurus* ARS-UCD2.0 reference genome which contains 21,667 protein-coding genes (Supplementary Data 2). Differential expression analysis revealed that in comparison with nRBC condition, 1654 genes were upregulated and 1618 were downregulated in S2P, meanwhile in R1A, 1496 were upregulated and 1769 were downregulated. The comparison between conditions S2P and R1A showed 700 upregulated genes in S2P relative to R1A, and 371 downregulated genes (Figure 1 A). A full list of differentially expressed transcripts for all three comparisons (S2P vs nRBC, R1A vs nRBC, and S2P vs R1A) is provided in Supplementary Data 2. Within protein-coding sequences, no substantial differences were observed in the number of differentially expressed genes (DEGs) between the S2P vs nRBC and R1A vs nRBC comparisons. Venn diagram analysis showed that 1866 transcripts (39.8 %) were differentially expressed in both S2P and R1A conditions vs nRBC, while showing no differential expression between S2P and R1A (Figure 1 B).

**Figure 1.**
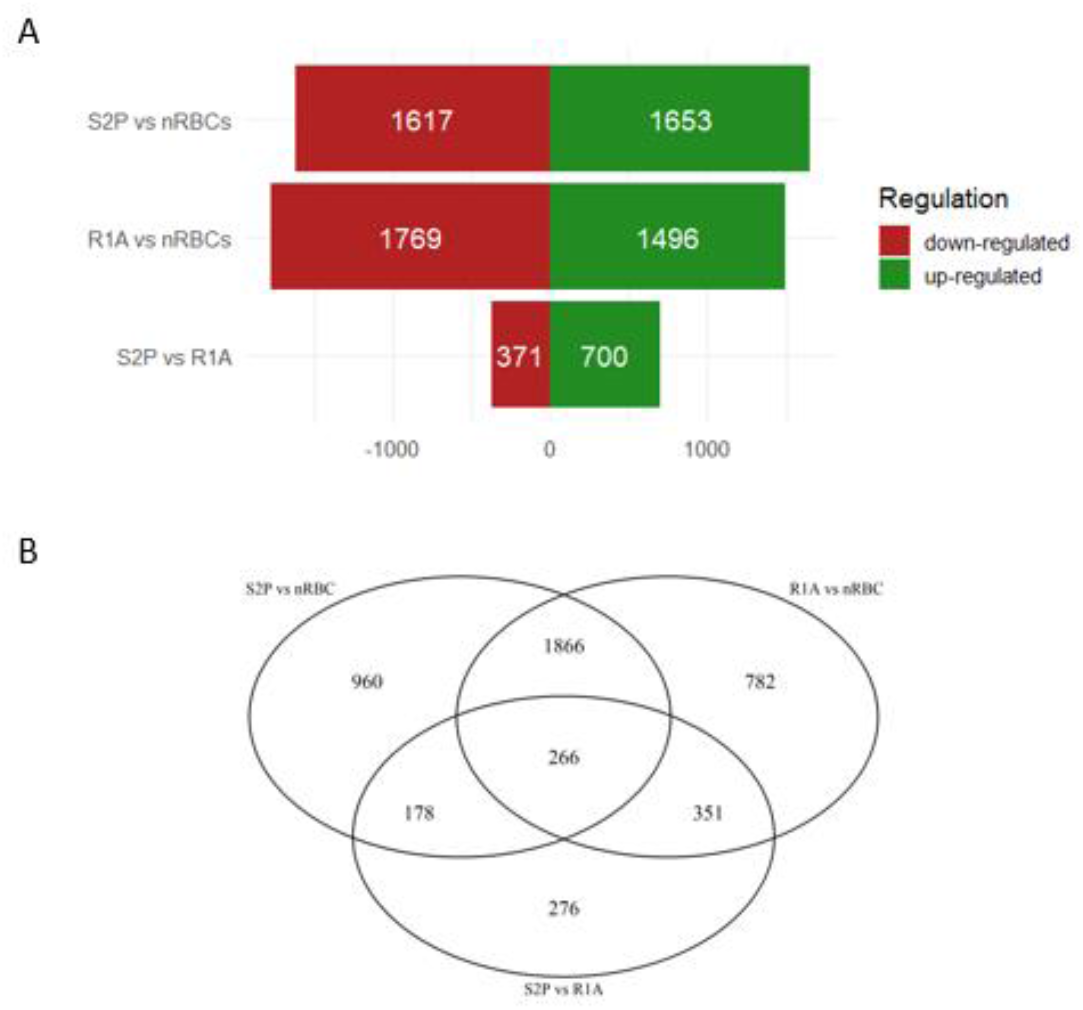
Differentially expressed genes in BMDMs co-cultures with *B. bovis*–infected erythrocytes or with nRBCs (A) Number of up- and downregulated genes identified in each experimental condition. (B) Venn diagrams of differentially expressed protein-coding transcripts illustrating shared and unique expression patterns between virulent and attenuated *B. bovis* strains.

### 3.3. Common host pathways modulated by virulent and attenuated *Babesia bovis* strains

Among the genes that were differentially expressed in macrophages upon phagocytosis of *B. bovis*–infected erythrocytes, regardless of whether the parasite was virulent or attenuated, we identified significant enrichment of common immune-related GO categories. These included terms associated with T cell differentiation and activation (e.g., GO:0045619, GO:0032946, GO:0050671), regulation of lymphocyte proliferation and migration (GO:0070665, GO:0045580, GO:0050871, GO:0042102), cytokine-mediated signaling pathways (GO:0045621, GO:0030888), and processes related to antigen processing and presentation (GO:0042130, GO:0031294, GO:0051251). Additional enriched categories involved regulation of adaptive immune responses (GO:0050864, GO:0032944), leukocyte-mediated cytotoxicity (GO:0050670, GO:0070663), and cell–cell adhesion during immune interactions (GO:0008645, GO:0042098, GO:0042129). Collectively, these results highlight that both virulent and attenuated *B. bovis* strains trigger a common transcriptional program in macrophages that is largely centered on immune-related pathways (Supplementary Data 3).

### 3.4. Strain-specific host responses to *Babesia bovis* infection

Among the 1071 genes differentially expressed between macrophages co-cultured with the virulent and the attenuated *B. bovis* strains, 700 genes were found upregulated and 371 downregulated (Figure 2).

**Figure 2:**
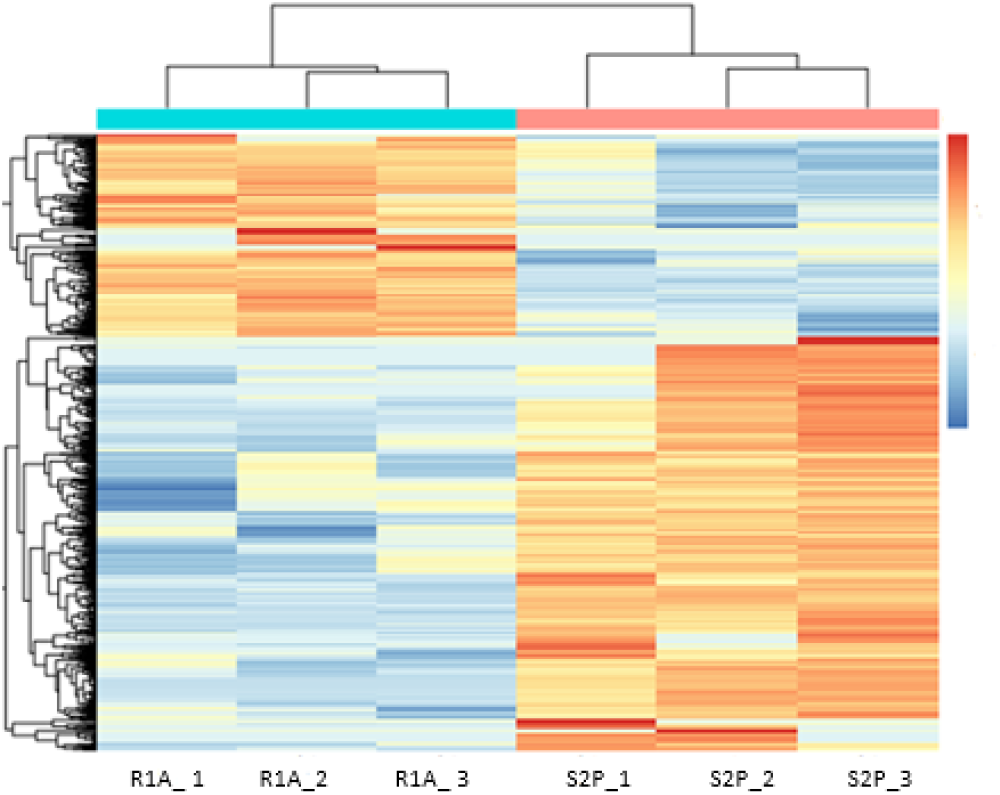
Heatmap illustrating DEGs between BMDMs that phagocytosed RBCs infected either the virulent strain (S2P_1, _2 and _3) or the attenuated strain (R1A _1, _2 and _3). The level of gene expression is represented by colors regarding log2FC: the higher score levels of expression are represented in red tones meanwhile the lower values are represented in blue tones.

Among the genes upregulated in the S2P condition compared to R1A, we found numerous genes associated with lymphocyte activation and adaptive immune responses, including PRF1, GNLY, ZAP70, IL2RB, RORC, ICOS, ITK, TBX21, GATA3, FOXP3, IRF4, STAT4, and CXCR3. Several co-stimulatory receptors and signaling molecules were also enriched, such as CD3D, CD3E, CD28, CD27, and CD6.

GO enrichment analysis highlighted terms related to lymphocyte and leukocyte activation, cytokine-mediated signaling, viral defense, and antigen receptor-mediated pathways (Figure 3 A; Supplementary Data 4). Moreover, processes such as response to type II interferon, negative regulation of viral replication, and hematopoiesis were significantly represented, underscoring a transcriptional program associated with immune response.

**Figure 3:**
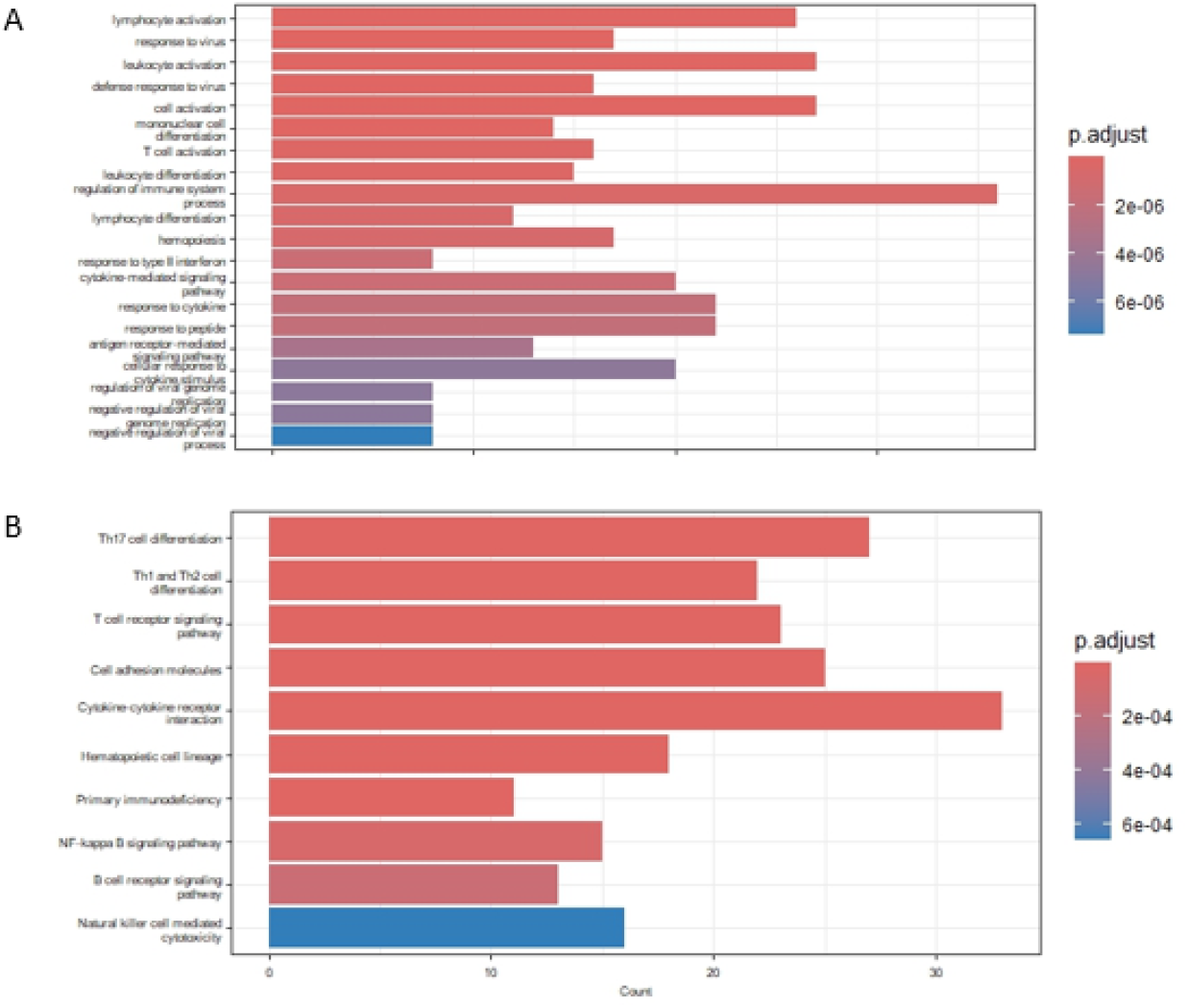
Functional enrichment analysis of genes up-regulated in BMDMs that phagocytosed erythrocytes infected with the virulent versus attenuated strain. A-Bar plot showing the top enriched GO Biological Process terms. B-Top enriched KEGG pathways. Bars represent the enrichment significance, measured as −log10 adjusted p-value.

Pathway analysis further supported this immune activation profile. KEGG analysis identified Th17, Th1, and Th2 enriched cell differentiation pathways, along with the T cell receptor signaling pathway, cytokine–cytokine receptor interactions, hematopoietic lineage commitment, and NF-κB signaling (Figure 3 B). Similarly, Reactome analysis identified pathways such as cytokine signaling, interleukin-4, interleukin-13, and interferon α/β signaling, and also RUNX1/FOXP3-mediated development of regulatory T cells. These results point out a coordinated activation of the effector and regulatory arms of adaptive immunity (Supplementary Data 4).

In contrast, a distinct transcriptional program characterized the genes downregulated in S2P vs R1A conditions. The downregulated genes included several regulators of cytokine and growth factor signaling (TGFBR1, VEGFC, IL18, IL1A, IL1B, IL23R, IL12RB2), chemokines (CXCL8, CXCL2, CXCL3, CXCL5, CXCL12), and structural and extracellular matrix components (COL1A1, COL1A2, COL3A1, COL6A1, COL6A2, LUM, SPARC). Notably, a high number of inflammatory mediators (S100A8, S100A9, S100A12, PTGS2, CCL2) and matrix metalloproteinases (MMP1, MMP2, MMP3, MMP12) were also significantly downregulated.

In line with this, GO analysis revealed a marked enrichment of processes related to chemotaxis, leukocyte and granulocyte migration, extracellular matrix organization, actin cytoskeleton dynamics, angiogenesis, and inflammatory response (Figure 4 A; Supplementary Data 5). These categories highlight a difference in gene expression of the pathways involved in cellular motility, tissue remodeling, and innate immune effector mechanisms when macrophages ingest the two *B. bovis* strains.

**Figure 4:**
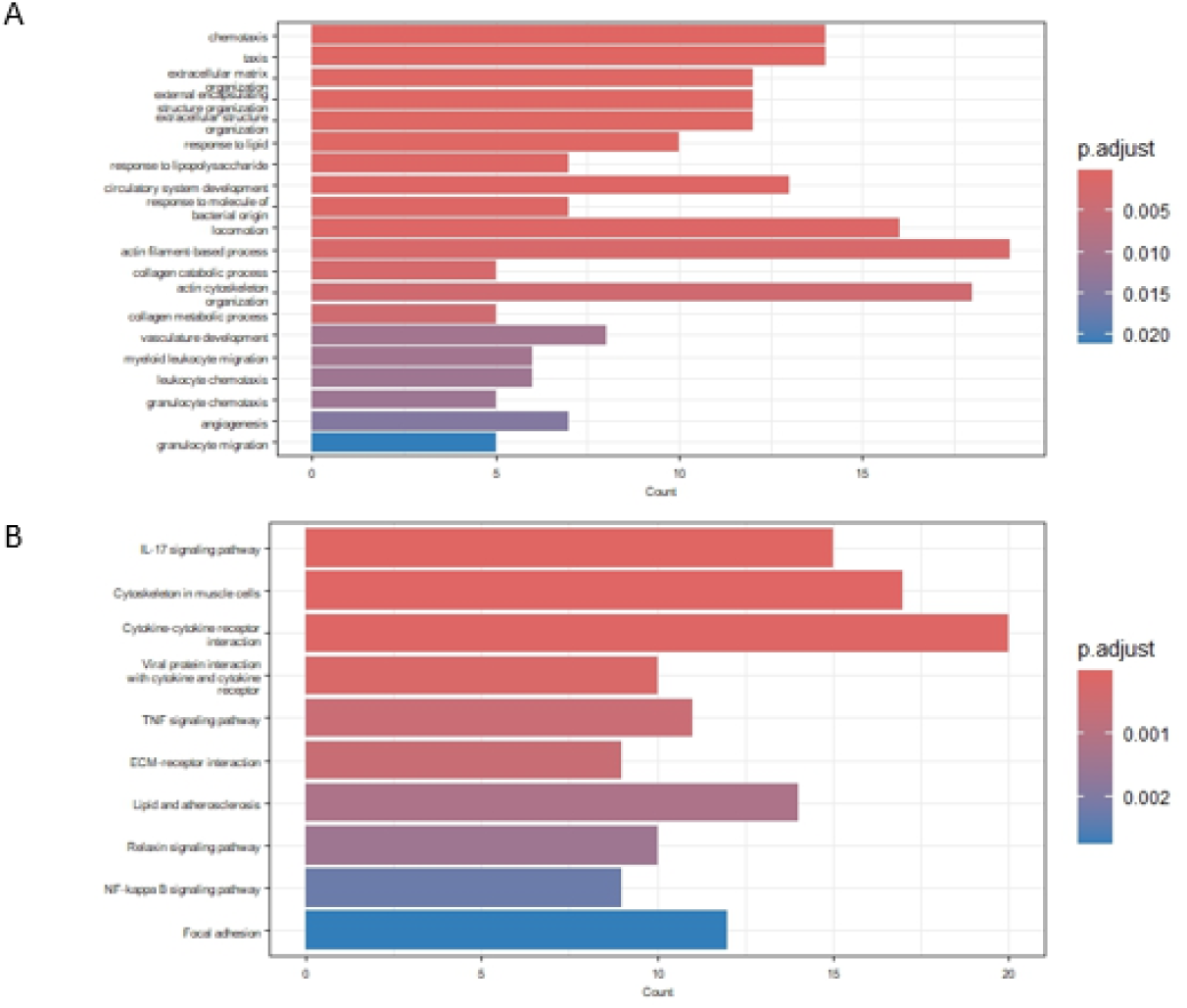
Functional enrichment of genes down-regulated in macrophages exposed to erythrocytes infected with the virulent versus attenuated strain A-Bar plot showing the top enriched Gene Ontology Biological Process terms. B-Top enriched KEGG pathways. Bars represent the enrichment significance, measured as −log10 adjusted p-value.

Pathway analysis reinforced this observation (Figure 4 B). KEGG enrichment showed significant association with the IL-17 and TNF signaling pathways, cytokine–cytokine receptor interactions, chemokine signaling, focal adhesion, ECM–receptor interaction, and NF-κB signaling, were downregulated in the virulent condition compared to the attenuated context, indicating that multiple pro-inflammatory and tissue remodeling networks were altered. Consistently, Reactome pathways results included extracellular matrix organization, collagen degradation, integrin interactions, interleukin signaling (IL-4, IL-10, IL-13), and NFE2L2-regulated antioxidant responses, suggesting broad host differences in extracellular remodeling, adhesion, and cytokine-mediated inflammatory signaling between exposure to the virulent and attenuated strains (Supplementary Data 5).

In summary, macrophages exposed to the two strains displayed distinct transcriptional programs, with the virulent S2P strain associated with enrichment of lymphocyte- and antiviral-related pathways. In contrast, the attenuated R1A strain induced a response characterized by stronger pro-inflammatory, chemotactic, and extracellular matrix remodeling signatures, highlighting clear differences in innate and adaptive immune engagement between the two strains.

## 4. Discussion

RNA sequencing has become a powerful approach to investigate host-pathogen interactions at the cellular level, since it enables the identification of specific pathways and molecular mechanisms that are modulated during infection, which would be difficult to uncover through targeted approaches. This strategy has been extensively applied to study infections caused by other apicomplexan parasites, including *Plasmodium falciparum* (Terkawi et al., 2018) and *Toxoplasma gondii* (Menard et al., 2021), providing valuable insights into how host immune cells detect and respond to parasite invasion.

In *Babesia bovis*, it is well established that attenuated strains can protect cattle against acute disease (Ristic, 2018). Given that common virulence genes were not identified among all virulent *B. bovis* strains or their attenuated derivatives (Lau et al., 2011) a plausible explanation is that they may generate distinct responses in the host, which in turn influence disease outcome. To test this hypothesis, we designed an experimental model that includes bovine macrophages as central mediators of the innate immune response against *Babesia bovis*. In natural infections, splenic macrophages recognize erythrocytes carrying damage-associated signals or surface proteins derived from the parasite and subsequently engulf them while presenting foreign antigens through MHC class II molecules. The *in vitro* model of BMDMs was successfully used by Valenzano et al., 2025 to identify novel *B. bovis* T-cell epitopes presented during infection, supporting its validity for dissecting host-parasite interactions. Although this interaction system may not perfectly mimic the behaviour of the bovine splenic macrophages, it is a feasible approach due to the lack of well-characterized continuous bovine macrophage cell lines.

The comparison between the three conditions allowed the identification of 4,679 DEGs out of the 21,667 protein-coding genes annotated in the *Bos taurus* genome (Liu et al., 2009). Among these, 1,866 genes were commonly differentially expressed in response to both the virulent and attenuated strains, indicating a core transcriptional response to *B. bovis* infection. Functional enrichment analysis revealed that these shared DEGs were significantly associated with immune-related GO categories. Collectively, these results indicate that both virulent and attenuated *B. bovis* strains elicit a conserved macrophage response strongly centered on lymphocyte activation, cytokine regulation, and adaptive immune signaling. The observation that most of the DEGs are related to immune responses, rather than to processes such as cell cycle regulation, proliferation, or migration, as has been reported for *T. gondii* (Menard et al., 2021), is consistent with the intrinsic life cycle of *B. bovis*, which only resides and multiplies in the red blood cell. For this reason, our results are consistent with the mechanisms employed by macrophages to process and present parasite antigens, which explains why the transcriptional response observed for both strains is predominantly immune-centered.

Among the immune-related genes reported in the natural response of bovines to infection by *B. bovis*, IFN-γ and TNF-α are particularly relevant molecules given their central roles in the Th1 response against the parasite (Brown et al., 2006; Goff et al., 1998). In our dataset both genes were downregulated when comparing both S2P and R1A conditions vs nRBCs, suggesting that infection with *B. bovis* may suppress their expression in macrophages. In line with this, IL-12 and NF-κB 1 and 2, were consistently reduced in both infections compared to control, which may contribute to reducing the transcription of pro-inflammatory mediators. The observation that the expression of these cytokines is reduced in BMDMs that phagocytised infected erythrocytes is consistent with previous findings in murine macrophages during infection with the related parasites of the genus *Plasmodium* (Wu et al., 2015). In this study, the authors showed that phagosomal acidification in macrophages suppresses the production of inflammatory cytokines, pointing out that dendritic cells constitute the major source of early proinflammatory mediators. Similar analysis with bovine dendritic cells co-incubated with *B. bovis* infected erythrocytes could shed light on whether this is a common trait between both apicomplexan parasites.

Another possible explanation for the lack of an inflammatory response in macrophages could be the absence of opsonization of the infected erythrocytes. Previous studies in *P. falciparum* have shown that opsonized infected RBCs stimulate human macrophages to secrete proinflammatory cytokines such as TNF and IL-1β, whereas unopsonized infected RBCs induce a much weaker inflammatory response (Ozarslan et al., 2019; Zhou et al., 2012). In the Babesia cultures used in this study, normal bovine serum from negative donors was employed, meaning that no parasite-specific antibodies were present in the co-cultivation system. Thus, this hypothesis is plausible and requires additional investigation.

The results described above were obtained after observing that one of the control replicates displayed a divergent expression profile, leading us to exclude it from further analyses. While several studies have demonstrated that reliable differential expression analyses can still be performed with two replicates when appropriate normalization and statistical methods are applied (Link et al., 2018; Srikumar et al., 2015), these results should be interpreted with caution. Importantly, the primary focus of this study was the comparison between the effects of virulent and attenuated strains on the host immune response, rather than contrasts with the uninfected control, which mitigates the impact of this exclusion on the overall conclusions.

The comparison of bovine macrophages exposed to erythrocytes infected with virulent versus attenuated *B. bovis*, showed clear differences in the expression of genes, highlighting lymphocyte- and antiviral-related pathways for the former and pro-inflammatory, chemotactic, and extracellular matrix remodeling signatures for the latter. Consistently, the higher number of upregulated genes in the virulent strain condition may suggest that the host response is stronger with that phenotype. A similar pattern has been reported for the apicomplexan parasite *T. gondii*, where infection with a highly virulent strain elicited a more pronounced transcriptional response in host cells than infection with a less virulent counterpart (Menard et al., 2021). However, a recent RNA-seq study of the bovine immune responses upon infection with *B. bigemina* showed that attenuated strain induces a stronger immune activation than the virulent one (Martínez-García et al., 2025). These apparent discrepancies can certainly be explained by differences in the host cell model (animal vs in vitro cell cultures) and the timing of the response assessed. While our study focuses on macrophages at an early stage of infection, the *B. bigemina* study analyzed peripheral blood mononuclear cells at the peak of disease, at day 8 after inoculation. This reinforces the importance of carefully comparing host responses across multiple cell types and infection stages, as they may reveal complementary aspects of the mechanisms driving virulence and attenuation, thereby adding significant value to the overall understanding of host-parasite interactions.

## 5. Conclusions

In this article, we investigated by RNA-seq the transcriptional responses in monocyte-derived macrophages that internalized RBCs infected with a virulent or an attenuated *B. bovis* strain. Aside from a common transcriptional response eliciting the upregulation or downregulation of several immune cellular processes, the different strain*s* also triggered a significantly different response in bovine macrophages, where several functional pathways appear activated or deactivated in response to the different phenotypes of the strains used. Collectively, these data improved our understanding of the early transcriptional events that occur in macrophages in response to *B. bovis*-infected RBCs and could be considered in further studies to better characterize the pathogenetic mechanisms of the disease as well as the comprehension of how live vaccines exert their effects.

## Acknowledgements

The authors would like to thank Lic. Pablo Vera for his assistance with library preparation and sequencing, and Dr. Julio Di Rienzo for his helpful suggestions on statistical analyses.

## Abbreviations

RBC: Red Blood Cell
IFN-γ: Interferon Gamma
TNF-α: Tumor Necrosis Factor Alpha
RNA-seq: RNA Sequencing
BMDM: Bone Marrow-Derived Macrophage
PBS: Phosphate-Buffered Saline
MOI: Multiplicity of Infection
DEPC: Diethyl Pyrocarbonate
FDR: False Discovery Rate
PCA: Principal Component Analysis
GO: Gene Ontology

## Supplementary Data captions

Supplementary Data 1: PCA of RNA-seq samples. Each point represents one replicate (n = 3) for each experimental condition. Axes indicate the percentage of variance explained by the first and second principal components. Clustering replicates reflects the similarity of gene expression profiles within each condition.

Supplementary Data 2: Lists of differentially expressed transcripts for all three pairwise comparisons: S2P vs uninfected, R1A vs uninfected, and S2P vs R1A.

Supplementary Data 3: Gene Ontology (Biological Process) enrichment analysis of DEGs shared between S2P and R1A conditions compared to uninfected nRBCs.

Supplementary Data 4: Functional enrichment of genes up-regulated S2P vs R1A A-Gene Ontology (Biological Process) enrichment analysis B-KEGG enrichment analysis C-Reactome analysis.

Supplementary Data 5: Functional enrichment of genes down-regulated S2P vs R1A A-Gene Ontology (Biological Process) enrichment analysis B-KEGG enrichment analysis C-Reactome analysis.

